# Computational identification of cell-specific variable regions in ChIP-seq data

**DOI:** 10.1101/796383

**Authors:** Tommaso Andreani, Steffen Albrecht, Jean-Fred Fontaine, Miguel A. Andrade-Navarro

## Abstract

Chromatin immunoprecipitation followed by sequencing (ChIP-seq) is used to identify genome-wide DNA regions bound by proteins. Given one ChIP-seq experiment with replicates, binding sites not observed in all the replicates will usually be interpreted as noise and discarded. However, the recent discovery of high-occupancy target (HOT) regions suggests that there are regions where binding of multiple transcription factors can be identified. To investigate ChIP-seq variability, we developed a reproducibility score and a method that identifies cell-specific variable regions in ChIP-seq data by integrating replicated ChIP-seq experiments for multiple protein targets on a particular cell type. Using our method, we found variable regions in human cell lines K562, GM12878, HepG2, MCF-7, and in mouse embryonic stem cells (mESCs). These variable-occupancy target regions (VOTs) are CG dinucleotide rich, and show enrichment at promoters and R-loops. They overlap significantly with HOT regions, but are not blacklisted regions producing non-specific binding ChIP-seq peaks. Furthermore, in mESCs, VOTs are conserved among placental species suggesting that they could have a function important for this taxon. Our method can be useful to point to such regions along the genome in a given cell type of interest, to improve the downstream interpretative analysis before follow up experiments.

## INTRODUCTION

A series of genome-wide experiments are largely adopted to study biological systems in relation to a given protein. They contribute to our understanding of particular molecular mechanisms at the basis of biological processes such as transcription and development, just to mention a few. In particular, ChIP-seq evaluates the genomic positions bound by a protein in the genome. Standard ChIP-seq experiments typically include replicated measurements in the experimental design in order to have the proper statistical power for the identification of reliable binding sites (or ChIP-seq peaks).

Previous results have indicated for several model organisms such as yeast, *Drosophila* and *Caenorhabditis elegans* the existence of genomic regions that are bound more often with respect to others, even in genomic positions in which a binding site is not expected for the protein under investigation. These regions have been previously characterized and described as “hyper-ChIPable” in yeast (1) and confirmed later in *Drosophila, C. elegans* and mouse and referred as “phantom peaks” (2). Furthermore, other regions defined here as variable regions, have protein binding that tends to variate stochastically and is difficult to interpret because their inconsistency in the reproducibility of the results. Current approaches to analyse ChIP-seq experiments do not report to the users regions that misbehave before downstream interpretative analysis; this might lead to the misinterpretation of the ChIP-seq results in terms of the function associated to the protein under investigation.

Here, we present a method that uses replicated ChIP-seq data for several proteins on the same cell line to detect regions that misbehave in ChIP-seq experiments. We assigned the term variable for a given genomic region if a protein binding site (or ChIP-seq peak) was not consistently detected in several experimental replicates of the same protein and for several independent proteins in a given cell type. These assignments can increase the value of ChIP-seq experiments by categorizing certain peaks as having cell-specific variability. Possible reasons for this variation might be the adoption of variable genomic structures (3), the high expression of a nearby gene (2), the specificity of the antibody used and the conformation of the chromatin during the immunoprecipitation. By finding variable regions, we expect to be able to characterize the origins of this variability and its potential relation to biological processes.

During the last years, the ENCODE consortium (4) addressed the problem of data collection for ChIP-seq experiments as well as other sequencing datasets creating the metadata of all the experiments. This effort is praiseworthy because at the time of reusing specific datasets it is important to know in detail how the data were produced, from which laboratory and according to which experimental criteria. This information allows controlling possible confounding factors in our study that focuses on local variability potentially caused by local genomic structural conformation or activity. Thus, we used data from the ENCODE consortium, and we controlled how the experiments were performed, from which laboratory and the bioinformatics tools used for data handling among other parameters.

In this work, we took advantage of the metadata provided by the ENCODE consortia, as indicated above, to select experiments in a consistent and comparable manner to implement a sliding window approach to classify genomic regions as variable or not.

Our results show that the method can identify variable regions (which we name variable occupancy target regions or VOTs) for every cell line tested and that, particularly for the K562 cell line, for which many datasets are currently available, it improves the separation of the samples in a PCA to promote a better downstream interpretative analysis. Method and scripts can be found online in this link: https://github.com/tAndreani/IPVARIABLE.

## MATERIAL AND METHODS

### Collection of ChIP-seq data

The ENCODE data portal provides comprehensive information about the meta-data of each experiment generated by the ENCODE consortium. We selected experiments according to specific parameters in order to avoid unwanted variability and to maintain consistency on the parameters of the downloaded data. The experiments were selected according to the following criteria: (i) laboratory producing the data as Snyder, (ii) identical untreated isogenic human cell lines (K562, MCF-7, GM12878, and HepG2) and ES-E14 mouse embryonic stem cells (mESCs), (iii) data processed with the standard ENCODE pipeline that uses the optimal IDR threshold as statistical method to obtain the significant peaks (5), (iv) status as released corresponding to a possible usage of the data, (v) the experiments of each biological replicate correspond to a peak file compared with appropriate input control experiment and (vi) peaks significance selected with a false discovery rate (FDR) lower than or equal to 5%, which results in a list of peaks with a fold change enrichment >= 2.25 in mESCs and >= 10 for the other cell lines. The metadata presented in JSON format was extracted and stored in a relational SQL database (See Supplementary Table S1 and Supplementary Fig. S1). For every cell we selected the following targets: for HepG2 we used MAFK, MNT, TBX3 and ZNF24 with two biological replicates; for MCF-7 we used CREB1, CLOCK, NFIB and ZNF512B with two biological replicates; for GM12878 we used BHLHE40, EP300, IKFZ2 and ZNF143 with two biological replicates; for K562 we used ARNT, NCOR1, MNT and ZNF24 with three biological replicates; for ES-14 mESCs we used HCFC1, MAFK, ZC3H11A and ZNF384 with two biological replicates (see Supplementary Table S1 for details).

### Reproducibility score implementation

After the identification of suitable experiments, the genome is binned in consecutive segments of 200 base pairs (bp) and the experimental ChIP-seq peaks are mapped to each segment. We formalized the reproducibility and not reproducibility of the segments for a given protein as illustrated in Fig. 1A and as follows:

**Figure 1.**
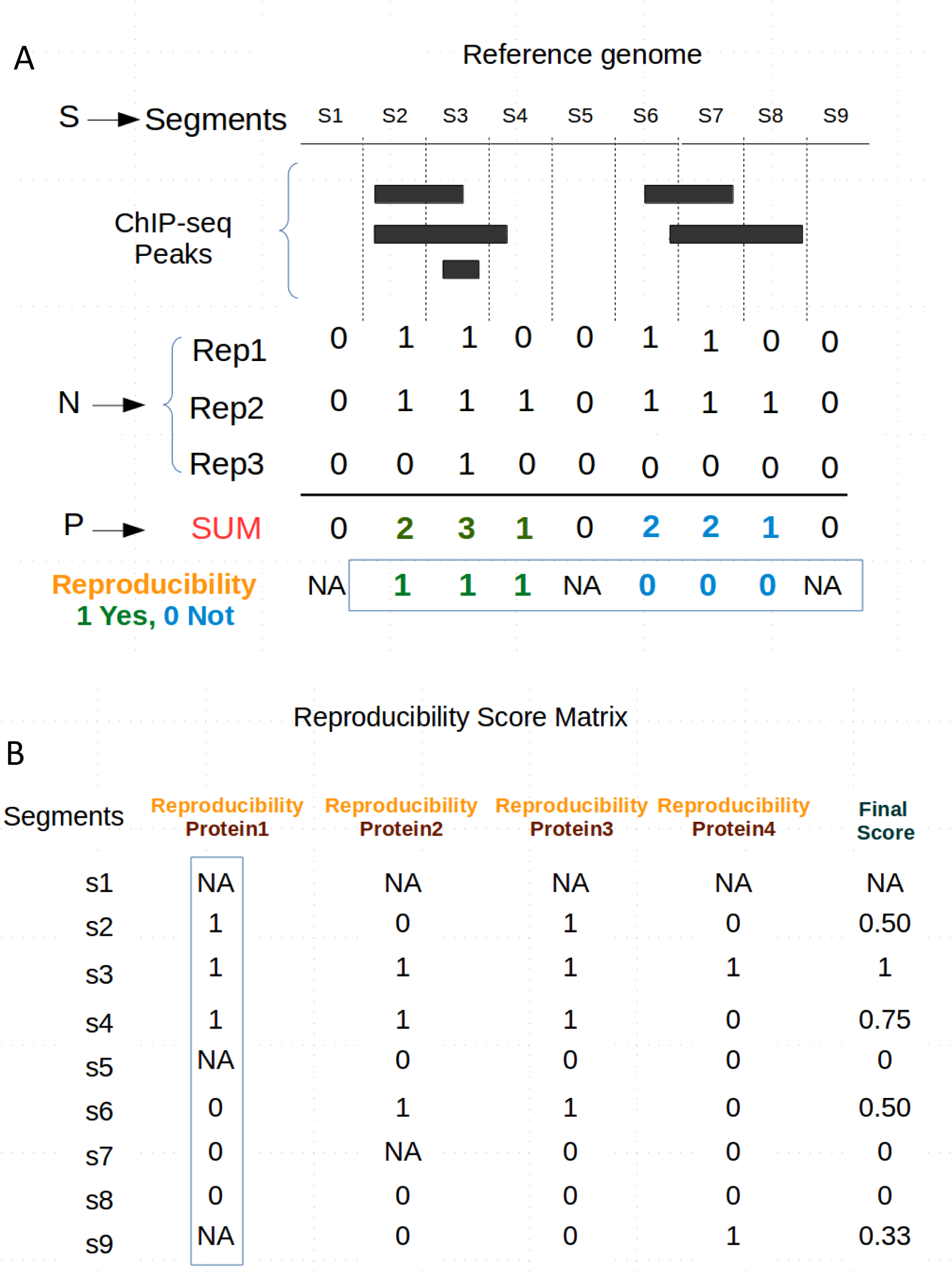
Method to annotate genomic regions with a reproducibility score. (**A**) From the genomic segments S, segments with ChIP-seq peaks for a given protein 1 in a given cell, in N=3 replicates, are converted to a binary format (Rep1 to Rep3). The sum at each segment of the values for the replicates (SUM or P) allows to define blocks of consecutive segments between zero-scored segments devoid of peaks (here two blocks; green and blue). All segments in a block are identified as indicating a reproducible region (Reproducibility=1; green) if the block holds at least one segment with value 3. Otherwise they are given a value indicating a non-reproducible region (Reproducibility=0; blue). (**B**) Average reproducibility values for ChIP-seq experiments from four different proteins in cell type A are combined in a final score that ranges from 0 (not reproduced in the four proteins) to 1 (reproduced in the four proteins). Only segments with values for at least 3 proteins were considered.

Let S be the genomic segments for a given genome;

Let N be the number of replicate ChIP-seq experiments for a given protein;

For each segment in S;

> Let P be the number of peaks detected in the segment;
>
> Reproducibility score = NA if P = 0;
>
> Else Reproducibility score = 1 if the segment itself or one of its neighbours* has P=N;
>
> Otherwise Reproducibility score = 0

* Neighbours are all consecutive segments with P > 0

In the following paragraph, we explain the procedure described by the pseudocode above in words. For our study, segments of the genome are defined considering a window size of 200 base pairs, N represents the number of replicates for each protein under investigation in a given cell type, and P is the number of replicated ChIP-seq peaks detected in a genomic segment (the signal). Consecutive segments without any signal (P=0; no peaks) are assigned with a NA. Consecutive segments in between two NA segments with a signal P reaching as a maximum value N are considered as reproducible regions and assigned a value of 1. On the contrary, consecutive segments in between two NA segments reaching a maximum value lower than N are considered as variable regions and assigned a value of 0 (Fig. 1A). The results of each protein under investigation are aggregated in a Reproducibility Score Matrix (RSM) (Fig. 1B) where rows show segments and columns show their reproducibility score for each protein and a final score (FS) defined as the average value of the row (or NA if more than 1 reproducibility score equals NA).

### Statistical test of scored regions

To assess whether the number of reproducible or variable regions associated with a particular score is significant, a suitable control had to be identified. The appropriate null distribution was built by randomizing the RSM. We performed this task using the “sample” function in R. The randomization was performed 1000 times, and regions at particular scores (0, 0.25, 0.33, 0.5, 0.66, 0.75 and 1) of the null distribution were counted. Afterwards, a *z-score* was computed according to this formula:

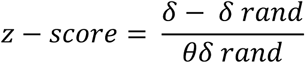

Where *δ* is the number of regions observed with a particular score, and *δ rand* and *θδ rand* are the mean value and the standard deviation of the null distribution, respectively. Assuming normality of the null distribution, it is possible to analytically calculate the corresponding *p-value* for a given *z-score* with significant level α = 0.05. The regions for each particular score were subjected to the test.

### Principal Component Analysis and Euclidean distances

Principal Component Analysis (PCA) was performed using the Python package scikit-learn version 0.19.1. The dots represented in the PCA are biological replicates for a given protein. Each colour represents a specific protein and the features set used to perform the PCA are all the segments detected in all the proteins. In order to test the effect of the removal of the variable regions in the PCA, segments within the variable regions were removed from the features set. The similarity distances between replicates of the same protein in the PCA were computed with the Python package SciPy version 0.19.1 using as a metric the Euclidean distance. Boxplot and Dotplot were performed using the Python library Matplotlib version 2.2.2.

### Enrichment analysis at regulatory elements

We collected genomic coordinates of the following gene related features from the UCSC table browser database in hg19 and mm10 annotations: promoter, 5’-UTR, coding exon, intron and 3’-UTR. Furthermore, we also used regions with R-loops (6) since they were previously reported as a potential feature associated with misbehaving ChIP peaks (3). For a set of regions (e.g. VOTs), the enrichment for each feature is obtained by dividing the number of regions overlapping a regulatory feature by the number of randomized regions overlapping the same feature.

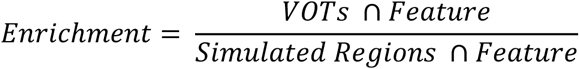

Randomized regions were obtained using bedtools version v2.25.0 shuffleBed (7).

### CG and AT dinucleotide frequency calculation

The percentage of CG and AT dinucleotides in the mouse and human genomes was calculated with the *nuc* function in the bedtools version v2.25.0 toolkit. To compute the CG and AT enrichment in the variable regions of mESCs and K562 cells we used a set of control regions using the shuffleBed function in bedtools version v2.25.0 (7). The differences in dinucleotide composition between the variable regions and the set of control regions were tested for significance using a t-test.

### Prediction of variable regions in K562 and mESCs using genomic features

In order to found out whether different genomic features (active chromatin marks, repressive chromatin marks, DNA accessible regions, CpG islands, etc.) could be used by a random forest classifier to predict variable regions in mESCs or in K562 cells we used a large panel of datasets.

For mouse ESCs, we also considered regions undergoing TET oxidation and bivalent domains. We used the following published data: 5hmC, 5fC and 5caC (8); CpG islands extracted from UCSC table browser for mm10 annotation; H3K4me1, H4K4me3, H3K79me2, H3K27ac, H3K27me3, H3K36me3 (9), LMR (10), DNAse-seq from ID:ENCSR000CMW experiment in the ENCODE portal. mm9 genome features were converted to mm10 using the Batch Coordinate Conversion (liftOver) tool from the UCSC Genome Browser Utilities (https://genome.ucsc.edu/cgi-bin/hgLiftOver).

For K562 cells, we used the following data downloaded from the ENCODE data portal: H3K27ac (ID: ENCFF044JNJ), H3K27me3 (ID: ENCFF145UOC), H3K4me1 (ID: ENCFF183UQD), H3K4me3 (ID: ENCFF261REY), H3K79me3 (ID: ENCFF350GQM), H3K36me3 (ID: ENCFF537EUG), DNAse-seq (ID: ENCFF856MFN). CpG islands were extracted from UCSC table browser for hg19 annotation.

To train and test the random forest model we used the function randomForest from the RandomForestClassifier from the Python package sklearn (0.21.3) (11). The default settings of the classifier were used except from *max_features* that was set to 2. As a positive set we have used the VOTs estimated by our method and as a negative set we used a set of regions obtained with the package gkmSVM version 2.0 (12). This package has a function named genNullSeqs capable of using the positive set of sequences and learning their nucleotide composition. Subsequently, the function generates a set of genomic locations with sequences of nucleotide composition and length similar to those in the positive test set. Since the genomic context is also important, we matched the simulated sequences within genomic features enriched in the VOTs such as CG rich promoters, 5’-UTRs and R-loops. The matching was performed maintaining the same number of regions found in the VOTs at each of those features. A similar approach was successfully used previously for the prediction of double strand breaks at CTCF and accessible chromatin sites (13). The random forest classification models were evaluated with receiver operating characteristic (ROC) curves within a stratified ten-fold cross-validation using the appropriate sklearn functions. In order to analyze the importance of single features contributing to the classification model RandomForestClassifier implements the function *feature_importances* that we used to receive values describing this aspect.

### Calculation of the evolutionary conservation of VOTs

To calculate the evolutionary conservation of VOTs we used the phastCons tool (14). The evolutionary score for each 200 bp segment was calculated by averaging the phastCons score of each nucleotide. As a null model, we used sequences in length, nucleotide composition and genomic context similar to VOTs. The test for significance was performed using the Wilcoxon rank-sum test with continuity correction. This operation was performed across 60 vertebrate species by using the phastCons60way tool in UCSC genome browser. Furthermore, this operation was performed across 60 placental vertebrates using phastCons60wayPlacental.

## RESULTS

### VOTs at transcription factor binding sites but not at chromatin marks can be detected in all tested cell lines

Our method to detect variable regions in ChIP-seq datasets for a given cell line relies on having several proteins tested for the particular cell line with replicates coming from the same laboratory and using the same platform (see Methods for details). Currently, the number of such sets in the ENCODE database is limited. While we were able to obtain a suitable set for K562 with four proteins and three replicates each, it is more usual to have a lower number of replicates, typically two.

For this reason, we applied our method to identify VOTs using just two replicates per protein for the human cell lines K562, GM12878, HepG2, MCF-7, and for mouse ESCs (mESCs). We found that K562 cells have a higher and significant number of variable regions for a total amount of 483 (p-val = 6.3e-103; Fig. 2A see Methods for details) whereas the other three human cell lines GM12878, HepG2, MCF-7 have similar lower but also significant numbers: 61, 76 and 62, respectively (p-val = 1.1e-07, 5.9e-6, or 4e-5, respectively) (Fig. 2C-D). Furthermore, we used another popular cell line used for developmental studies, mouse embryonic stem cells ES-14. Also in this cell line, we identified a number of VOTs for a total amount of 332 (p-val = 6.9e-3) (Fig 2E).

**Figure 2.**
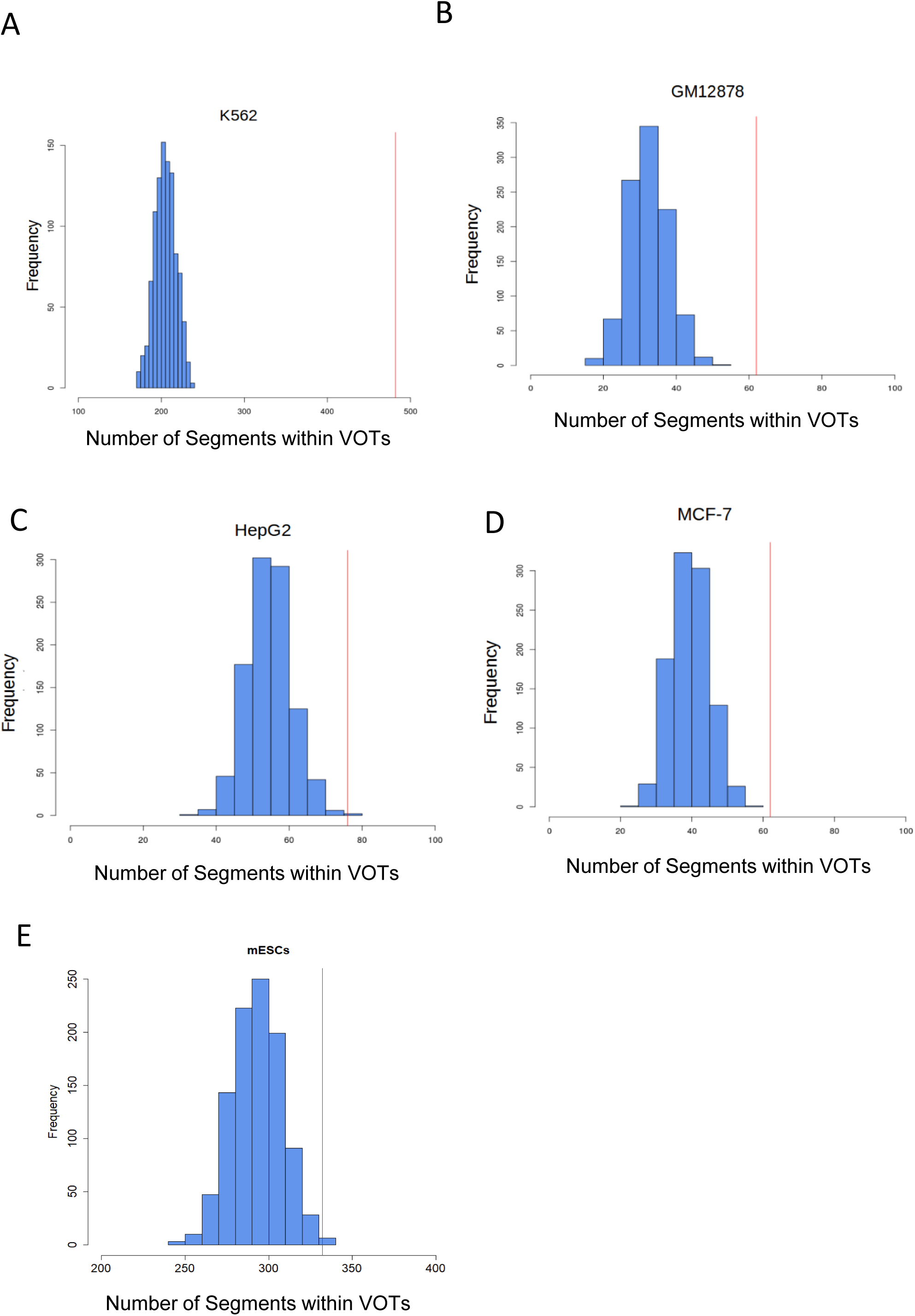
Significance of the observed number of variable regions for different cell types. The number of VOTs observed in each cell line (red line) is significantly higher than the corresponding computed null distribution (blue). (**A**) K562, (**B**) GM12878, (**C**) HepG2, (**D**) MCF-7 and (**E**) mouse ESCs cells.

We next asked whether the variable regions are also present at chromatin marks. For this, we have applied our method to K562, human ESCs (hESCs) and MCF-7. We found very few variable regions in K562 and hESCs for an amount of 8 and 7 respectively and given the low number we did not subject those regions to a statistical test. For MCF-7 we did not find any regions while for the other cell lines HepG2 and GM12878 there were not enough data to run our method. This led us to conclude that VOTs are typical of transcription factors and not of chromatin marks.

Finally, we wanted to see to which extent increasing the number of replicates in the experimental design would increase the number of variable regions. We were able to design a proper experiment only for K562 cell lines, which is the cell line with the higher number of experiments in the ENCODE project. Using three replicates per protein and four proteins (see Methods for details), the number of variable regions detected was drastically higher (a total of 3012). We also used the three replicates from this cell line to evaluate the reproducibility of the identified VOTs. Comparing replicate 1 vs. 2, 1 vs. 3, and 2 vs. 3, we obtain 486, 254 and 176 regions, respectively. The overlaps between the sets obtained are not random. VOTs from 1 vs. 2 overlapping with VOTs from 1 vs. 3 are 56 (p-value = 1.59e-16, Fisher test). VOTs from 1 vs. 2 overlapping with VOTs from 2 vs. 3 are 3 (p-value = 0.19, Fisher test). VOTs from 1 vs. 3 overlapping with VOTs from 2 vs. 3 are 14 (p-value = 1e-4, Fisher test).

### Variable regions are rich in CG dinucleotides and enriched along gene body features

Next, we tested the CG dinucleotide frequency of the variable regions in K562 and mESCs. We found a higher frequency of CG dinucleotides compared to a random set of control genomic regions in K562 (p val 3.8e-4) and mESCs (p val 8.4e-8) (Fig. 3A and 3B, respectively). The protein targets in the ChIP-seq experiments used do not have DNA-binding motifs particularly affine for CG dinucleotides (motifs from the Jaspar database (15); Supplementary Fig. S2), hence the CG composition is specifically related to the variable behavior and not to the DNA-motifs bound by the proteins selected.

**Figure 3.**
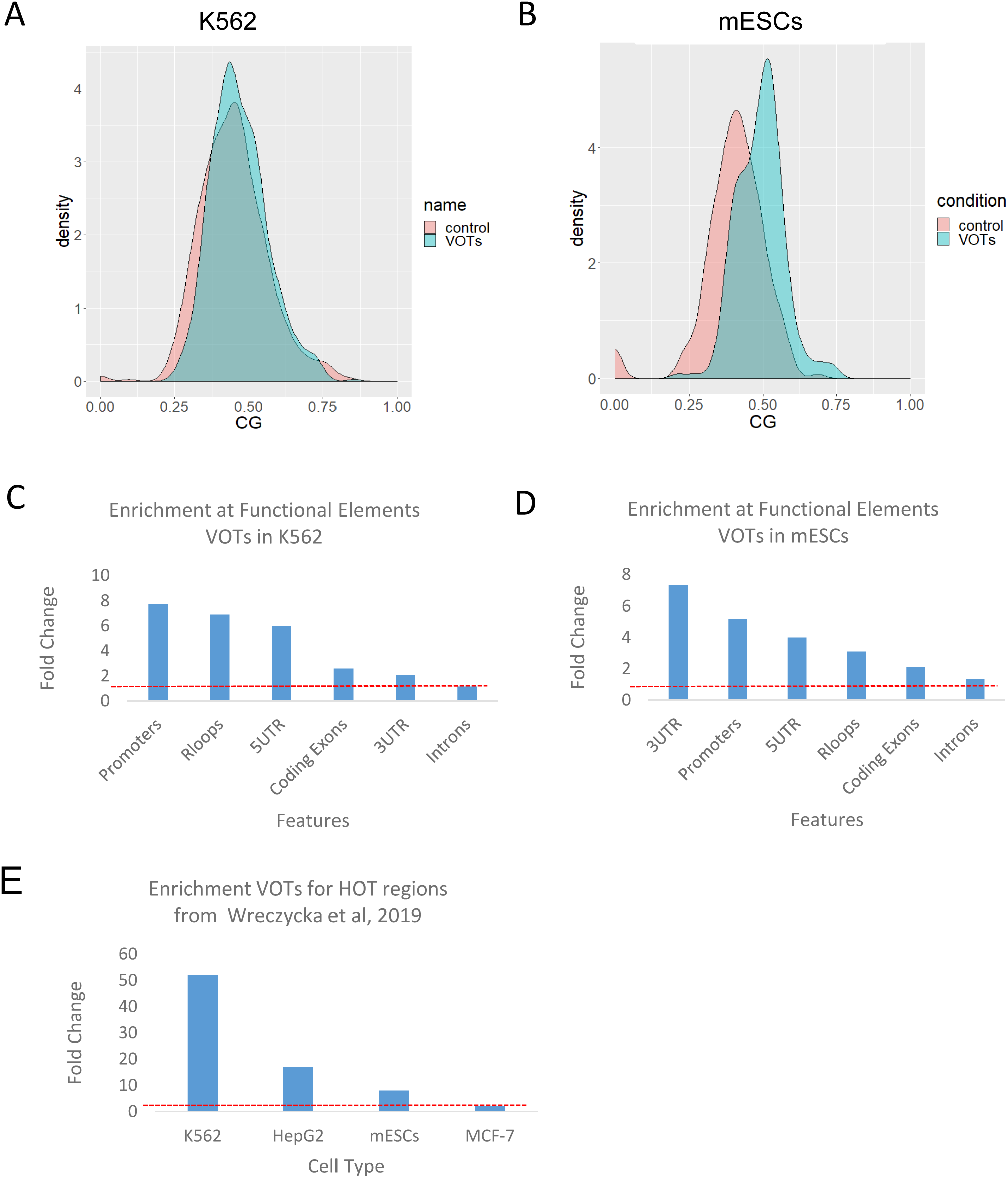
Properties of variable regions in K562 and mESCs. CG dinucleotide composition in K562 human cell lines (**A**) and in mouse ESCs (**B**). Control regions are a set of randomly sampled genomic regions of similar size. Enrichment of VOTs in K562 human cell lines (**C**) and mouse ESCs (**D**) at several gene body features and R-loops. (**E**) Enrichment of VOTs at HOT regions (3).

Furthermore, variable regions were highly enriched for different features among K562 and mESCs cells with K562 showing a high enrichment for promoters and R-loops (Fig. 3C) and mESCs showing a high enrichment for 3’-UTR and Promoters (Fig. 3D). In a recently published work (3), the authors reported that previously characterized regions as “high occupancy target” (HOT) (16,17) are likely to be a ChIP-seq artifact. Among the properties of these regions, they reported GC/CpG-rich kmers and RNA–DNA hybrids (R-loops). Since also in our work we found these characteristics for the variable regions, we downloaded the regions reported in (3) and checked for a possible enrichment. We found significant enrichment for all the cell lines tested except for GM12878. Especially for the K562 and HepG2 cell lines the enrichment was of 52 and 17 fold change, respectively (observed vs expected, two-sided Fisher test p val. 1.46e-87, 3.9e-3, respectively). Furthermore, also mESCs showed a significant enrichment of 8 fold change (observed vs expected, two-sided Fisher test p val. 3.2e-3) (Fig. 3E).

### Variable regions are not blacklisted regions from ENCODE project nor associated with structural variations in cancer cell lines

The ENCODE consortia provides a detailed description about the meaning of “blacklisted sites”. These genomic positions often produce artifact signal in certain loci mainly because of excessive unstructured anomalous sequences. Reads mapping to them are uniquely mappable so simple mappability filters do not remove them. These regions are often found at specific types of repeats such as centromeres, telomeres and satellite repeats. Given the high variability of the ChIP-seq peaks of the regions described in this manuscript we thought to check whether our method was detecting the already described and characterized “blacklisted regions” or not. To answer this question, we analysed the overlap of the variable regions obtained in all the cell lines we have used (K562, HepG2, MCF-7, GM12878 and mESC) with the public available ENCODE blacklisted regions (18). We found no overlap except for K562 (significant depletion, 35 observed vs 261 in random model, two-sided fisher test p val. 2.92e-46) and mESC (significant depletion, 7 observed vs 29 random, p val. 2.1 e-4). These results confirm that our variable regions are not associated with the ENCODE blacklisted regions, hence need to be considered for new detection methods. Finally, given the presence of structural variation in cancer cell lines, we asked whether VOTs found in cancer cell lines could be the outcome of such variations. To test this, we downloaded annotated structural variants from the https://portals.broadinstitute.org/ccle/ website for K562, MCF-7 and HepG2 and found no overlap for any of the corresponding VOTs. There were no data available for GM12878.

### The removal of variable regions improves the interpretation of the PCA in K562 cell lines

Variable regions may reflect cell-specific effects that are not target-specific. While this information might be indicating biological function, we hypothesized that the removal of such target non-specific data could result in an improvement of the separation of the replicates points in a PCA. In order to test such potential benefit in removing the variable regions for downstream interpretative analysis, we performed a PCA of the ChIP-seq samples obtained for the K562 cell line (Fig. 4A). The PCA was performed using (i) all the segments bound by each protein in the respective replicates in the original datasets and (ii) without the segments within the variable regions. We found that the separation along the components improves after removing the segments within the variable regions. Furthermore, the replicates of the proteins tend to cluster better without the segments within the variable regions and this is reflected with lower Euclidean distances in pairwise comparisons between replicates of the same proteins (Supplementary Fig. S3). We note that this does not mean that data from these regions should be discarded, but that they should be considered differently. Further research is needed to characterize these regions and find out if they have a cell-specific biological function.

**Figure 4.**
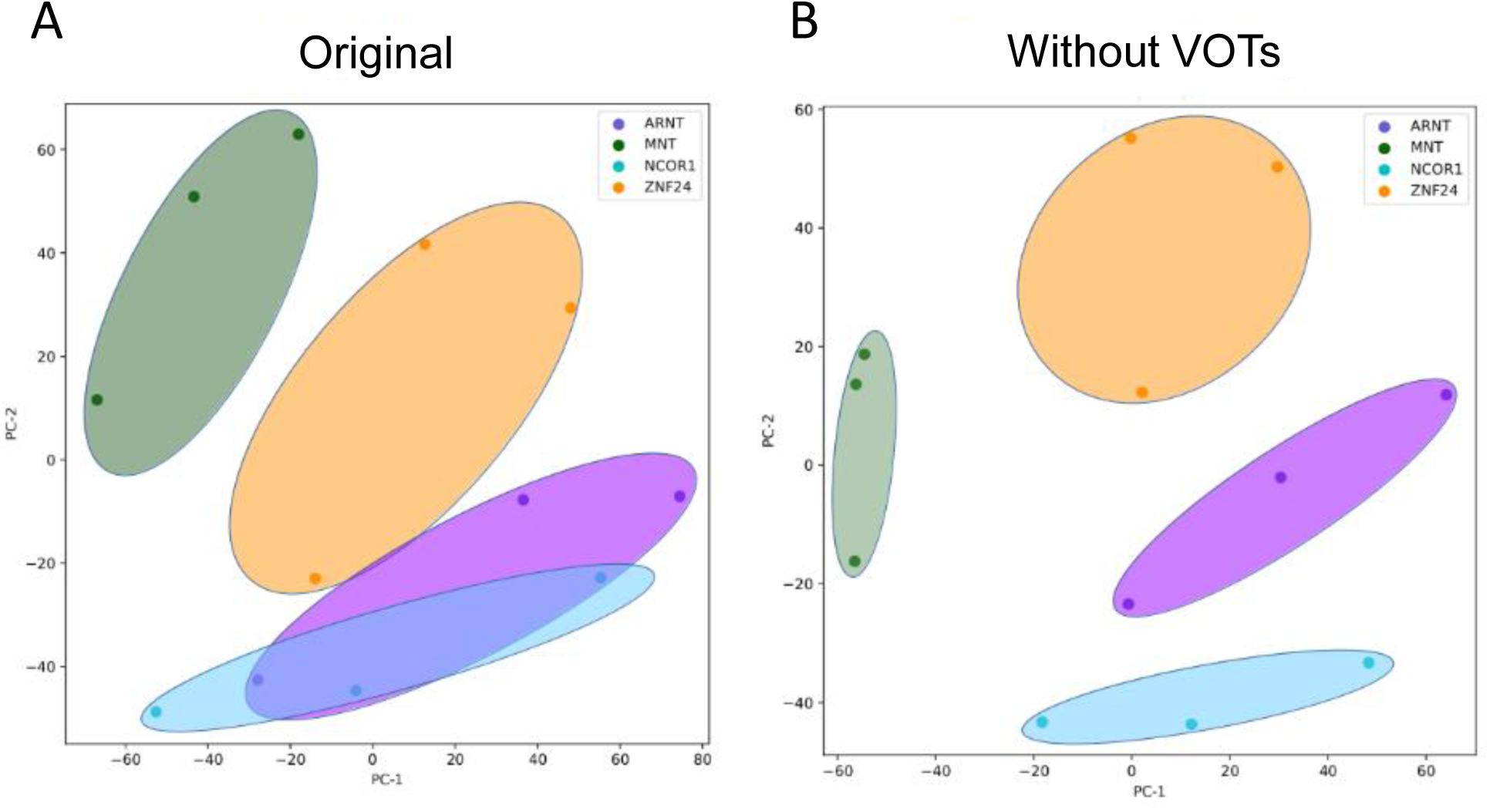
Effect of removing variable regions. PCA using presence of peaks in the set of 200 bp segments on the genome as features with the original data and without 200 bp segments within the variable regions in K562 cell lines (**A** and **B**, respectively). Each dot represents a biological replicate and each colour the protein target.

### VOTs are predictable by DNA accessible regions in K562 and mESCs

Finally, we searched for genomic features that can be predictive of variable behaviour. We evaluated the possible association of different genomic features with variable regions using a random forest classifier.

The classifier was trained with a positive set consisting of the variable regions detected in mESCs (332 variable regions), and with a negative set consisting of genomic sequences with size and nucleotide composition similar to those of the positive set (see Methods for details about the training and about the set of genomic features).

The algorithm was able to classify the variable regions (AUC (Area Under ROC-curve) = 0.718) and returned as best predictors DNA accessible regions, together with regions lowly methylated and oxidative products of TET enzymes 5hmC and 5caC (Fig. 5A and 5B). These modifications are highly frequent at distal regulatory elements (8) and promoters (19) and we speculate that the turnover of these modifications might affect the binding of the proteins to the DNA leading to stochastic variation of the binding sites.

**Figure 5.**
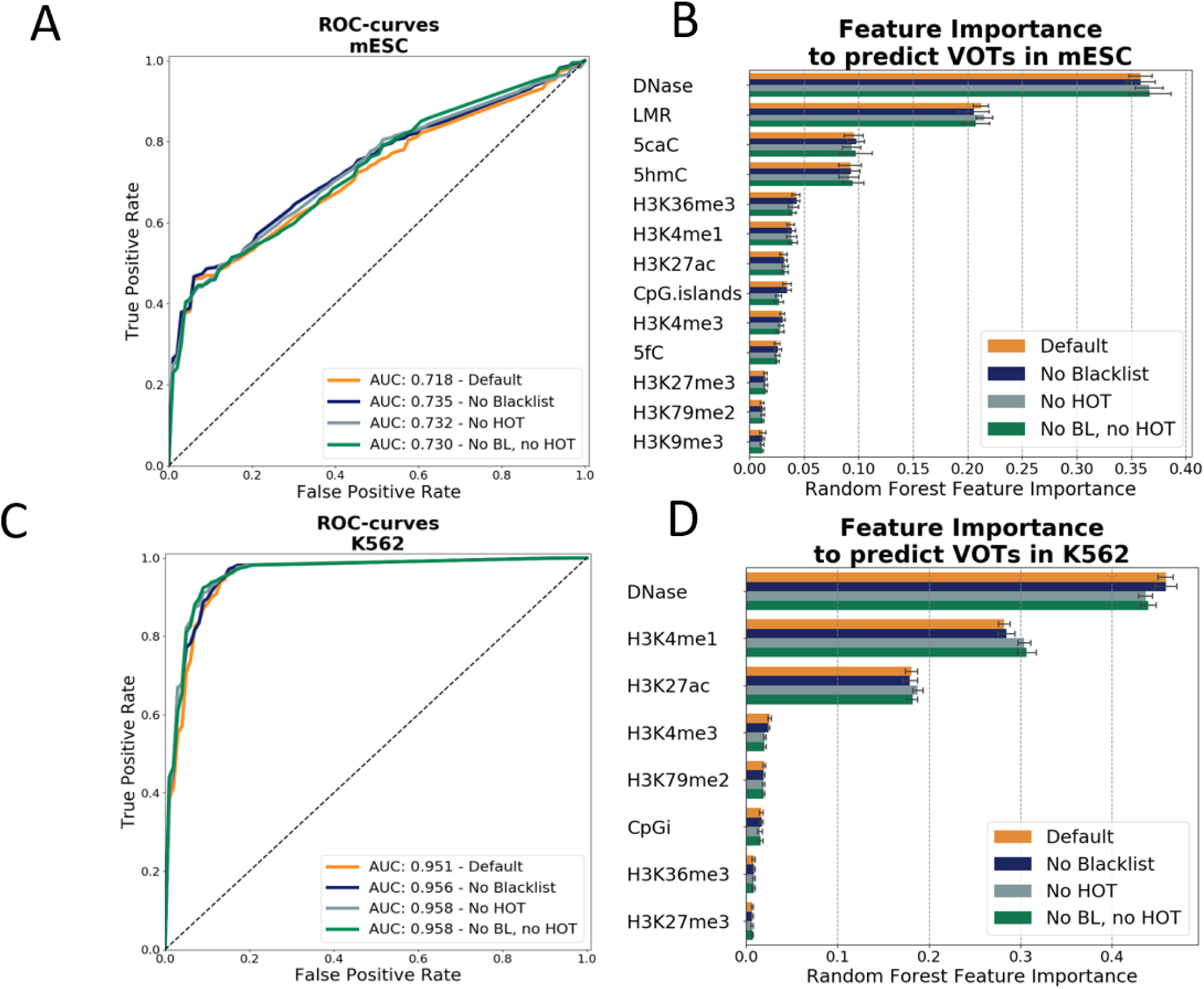
Prediction of variable regions in K562 and mESCs. (**A**) Receiver operating characteristic (ROC) as a quality measure of the predictability of the variable regions in mESCs and (**B**) importance of the features for predicting the variable regions in mESC measured as mean decrease Gini in the random forest. See text for details about every feature. (**C**) Receiver operating characteristic (ROC) as a quality measure of the predictability of the variable regions in K562 cells and (**D**) importance of the features for predicting the variable regions in K562 cells measured as mean decrease Gini in the random forest.

To observe the reproducibility of these results, we studied next data from K562 cells (483 variable regions). Again, as best predictor of variable regions we found DNA accessible regions, together with K3K4me1 and H3K27ac chromatin marks (AUC = 0.951) (Fig. 5C and 5D).

### Functional analysis and evolutionary conservation of VOTs

In order to study the functional implications of VOTs, we decided to interrogate Cistrome-GO (20) and perform functional enrichment analysis for K562 and mESCs, which were the two cell types with a reasonable number of regions for such an analysis. We found in K562 as a top hit the RNA Pol-II core complex (corrected p-value 1.72e-06, Fig. 6) suggesting that the variable regions can be located at sites of genes regulating core complexes of transcription. Consistently, for mESCs the only significant Gene Ontology term found was protein DNA complex (corrected p-value 3.3 e-02).

**Figure 6.**
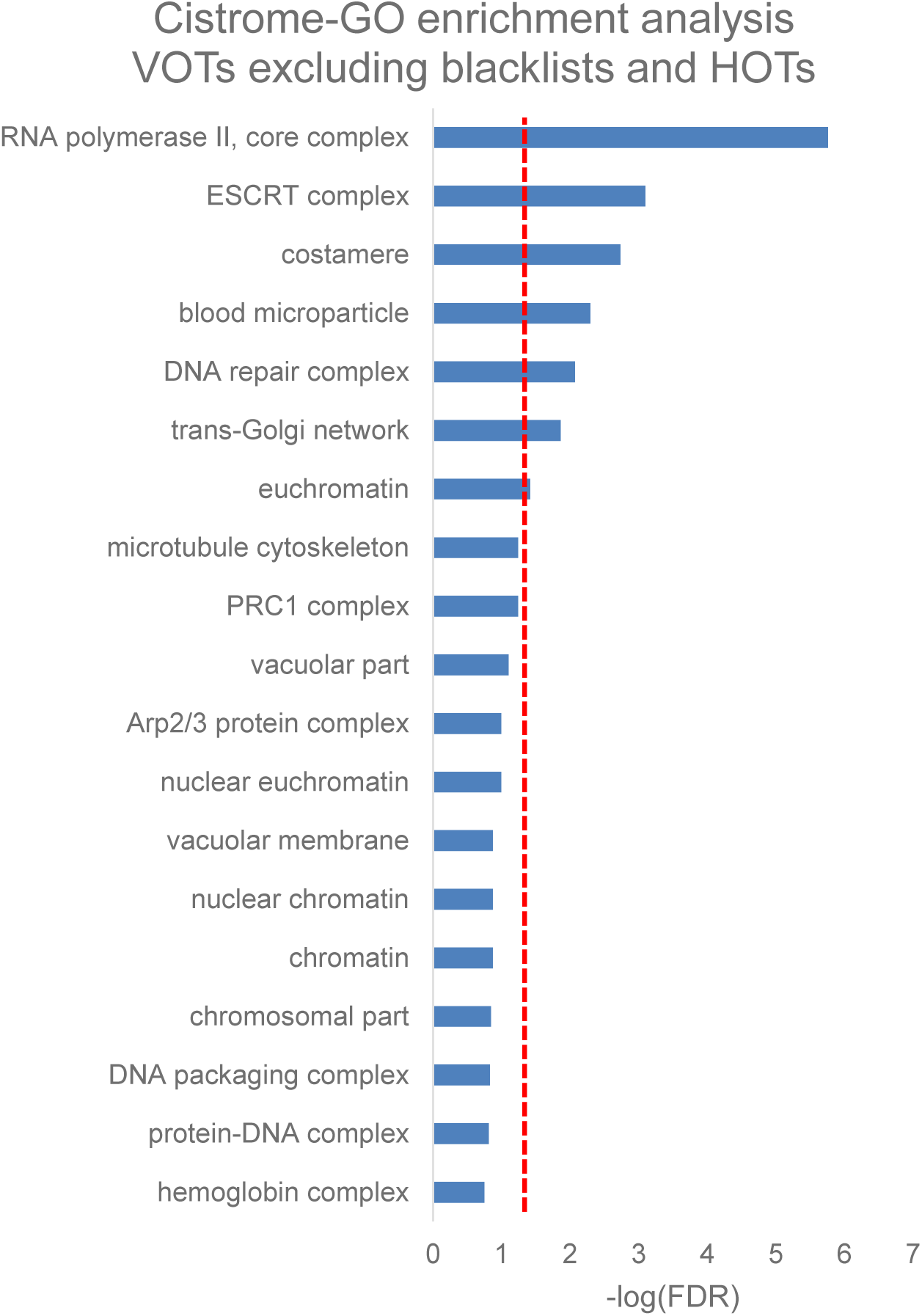
Functional analysis of VOTs. GO enrichment analysis of VOTs (not overlapping HOTs or blacklisted regions) for K562 cell lines using Cistrome-GO. A similar enrichment analysis for mESC selected only the term “protein-DNA complex” (see text for details). The dashed vertical red line represents FDR = 5%.

Considering that VOTs in mESCs could have a functional implication in development, we asked whether there is a conserved role for these DNA sequences across different vertebrate species. We performed this using the phastCons (22) tool (see Methods for details) and found that VOTs in mESCs are not conserved among 60 vertebrate species (Wilcoxon rank-sum test with continuity correction p-value = 0.12, Fig. 7A). However, considering only the placental among the 60 vertebrate species we found a significant result (Wilcoxon rank-sum test with continuity correction p-value = 0.021, Fig. 7B). This suggests that VOTs would have a function specific to this taxa.

**Figure 7.**
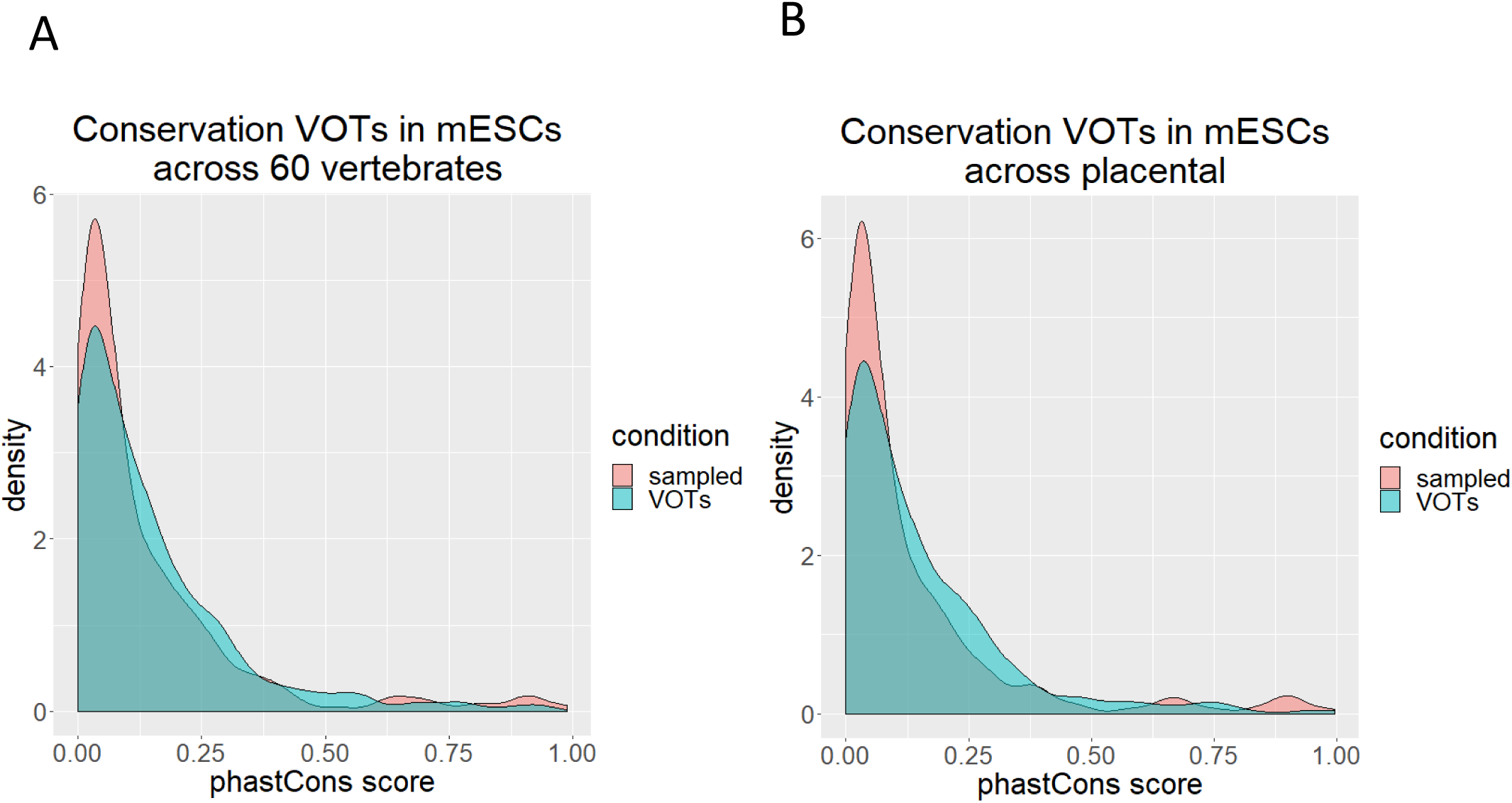
Conservation of VOTs in mESCs. (**A**) across 60 vertebrate’s species and (**B**) across placental. x-axis represents the average phastCons score in segments of 200 bp and y-axis represents the number of segments for the particular score.

Taken together, our results suggest that VOTs have a conserved role in the establishment of feedback loops of the gene regulatory network stochastically influencing the expression of DNA binding proteins at DNA accessible regions.

## DISCUSSION

During the last years, several laboratories tried to study the regulation of gene expression in different model organisms. For this scope, ChIP-seq was adopted as a standard technique but the extent of its usage raised some questions in terms of reliability (1,2,21,22). In particular, in a previous work (3), Wreczycka and colleagues presented a method that considers the nature of phantom peaks and hyper-ChIPable regions to define high-occupancy target (HOT) regions where un-specific binding to multiple targets would be found even in the absence of expected binding motifs. They concluded that the un-specificity of binding sites in HOT regions is associated with CG dinucleotide rich regions and as a consequence at R-loops (that are CG rich) and DNA tertiary structures. Though, this is a common concern for ChIP-seq assays and since the beginning the technique was known to be biased toward GC-rich contents during fragment selection in the steps of the library preparation and amplification during the sequencing (21). Here we have found evidence that supports that such regions could also be responsible for variable behaviour in ChIP-Seq in a cell-specific fashion. Our method evaluates replicated ChIP-seq experiments for multiple targets in a cell type, to find regions where target binding is not reproduced in all replicates for multiple targets. These variable-occupancy target regions (VOTs) are cell-specific and share structural features with HOT regions. However, differently to HOTs, VOTs do not produce consistent un-specific target binding. Accordingly, VOTs do not overlap blacklisted ENCODE regions. Together, the cell-specificity of VOTs and our finding that VOTs can be predicted using DNA accessibility suggest their dependency on gene expression and epigenetic state.

While the most consistent enrichment of variable regions observed was for promoters and 5’-UTR regions in both K562 cells and mESCs, the differences observed for variable regions at R-loops suggest that it is not possible to drive a certain conclusion relating where exactly this variation occurs. On the other hand, the fact that we are able to predict variable regions using genomic features alone with relatively high accuracy indicates that there is certainly a relation between genomic features and variability that could be eventually detected. Taking these results together, we assume that with the future availability of further ChIP-seq datasets testing multiple proteins in the same cell lines it will be possible to assess the sources of variability in ChIP-seq with more certainty. Regarding the predictive genomic features, we note that histone marks were not predictive of VOTs in K562 and mESCs. This is in agreement with the fact that VOTs are not detected when using ChIP-seq testing histone modifications (as opposed to ChIP-seq testing protein binding). These results support that VOTs are not due to ChIP-seq technical issues.

With our method, we propose a systematic approach using ChIP-seq experiments and replicated measurements in a given cell-type to identify misbehaving DNA regions that have to be treated differently in the post processing downstream analysis. We have shown that discarding data from these regions can improve studies focusing on target specific effects. However, the specific study of VOTs is necessary, since our functional analyses indicate that VOTs might regulate DNA binding proteins at regions enriched in R-loops and 5’-UTRs, which have a tendency to adopt structures. For mESC we found their conservation among placentals. These results suggest their role in developmental functions by feedback loops of the gene regulatory network. Formation of structures in VOTs could result in sites in which different transcription factors compete for binding in a stochastic manner that results in the observed variability. Further research is needed to study potential cell-specific functions of VOTs as hubs or sponges for transcriptional regulatory complexes, which could be verified with other experimental assays like ChIP-qPCR. We suggest applying our approach as a post processing quality check of the data before starting follow up experiments and driving biological conclusions.

We must point out that our method requires enough replicates for the same protein in a given tissue. For example, predictions for organisms like the fly *Drosophila melanogaster* using data from modENCODE and modERN are not yet possible, At this point, the only datasets we find suitable for our analyses are in ENCODE.

Finally, we note that similar approaches to the one used for our method to point to variable regions in ChIP-seq datasets could be eventually developed and applied to any type of next generation sequencing datasets that uses replicated measurements under various conditions (for example, ATAC-seq from multiple cell types). This could open avenues for the discovery of other types of variability leading to a more informed use of sequencing-based data. The study of the similarities and differences between variable regions obtained with different techniques might be crucial to increase our understanding of the inter-relation between genomic structural flexibility and regulatory function.

## DATA AVAILABILITY

The code to reproduce this manuscript is available in the GitHub repository (https://github.com/tAndreani/IPVARIABLE). The list of ENCODE experiments used is available as Supplementary Table S1. The coordinates of VOTs found in K562, HepG2, MCF-7, GM12878 and mESC are provided as Supplementary Table S2.

## SUPPLEMENTARY DATA

Supplementary Data are available online. **Supplementary Table S1.** Table with the list of ENCODE experiments used. **Supplementary Table S2.** Coordinates of VOTs found in K562, HepG2, MCF-7, GM12878 and mESC.

## ACKNOWLEDGEMENTS

We thank Katerina Taškova for fruitful discussions. Tommaso Andreani and Steffen Albrecht were affiliated with and financially supported by the International PhD Programme on *Gene Regulation, Epigenetics and Genome Stability*, Mainz, Germany.

## SUPPLEMENTARY MATERIAL

### Supplementary Material

#### Conversion of JSON-based files to a relational SQL database

The metadata in ENCODE is represented in JSON-format. There is an API (Application Program Interface) that enables downloading JSON files for many experiments automatically. In addition, it is possible to restrict the experiments to be downloaded by specifying conditions the experimental metadata has to comply with. However, a JSON file for one experiment can include up to 11,000 lines and this can cause two problems. The first is that the extraction of information can be complicated and requires further implementation of small scripts / programs. The second is that parsing such files can be computationally expensive especially if there are more than 1000 files to be processed. To avoid these problems and enabling the analysis of the metadata in a flexible and fast way we decided to convert the information from JSON into a MySQL database. For objects like experiment, target, organism, replicate, etc. we downloaded all JSON files. A Python script was implemented that extracts information encoded in each JSON file and automatically fills the SQL database with this information. The procedure in the Python script takes also cares about creating the tables for each object and furthermore creating also relational tables for the connection between two tables (objects). Relational information is also automatically extracted and stored into SQL. Figure 1 represents a simplified case with one experiment. The brackets “{}” define the beginning and end of a JSON. There are simple features like accession and status for which the value is directly given. Those simple values can also appear in lists represented by the brackets “[]”. However, one JSON can include other JSONs. This is shown by the target in the experiment that is completely described by a so called nested JSON. It is also possible that one JSON includes a list of JSONs. The example in Figure <u>S</u>1 shows that for each file that is related to the experiment there is one JSON within the experiment. These nested JSONs are the reason why those files are getting so large. Another advantage coming with the conversion from JSON into SQL is that objects are not stored redundantly. Taking as example the target that describes the protein that is targeted in a ChIP-seq experiment. All JSONs for experiments with the same target store the same information for the same target multiple times. In SQL there is only one entry for a specific target in the target table and experiments with this target are just linked to this entry by the target_ID. An example for such a link is represented by the target in Supplementary Figure S1 below and has_file is a relational table.

**Figure S1.**
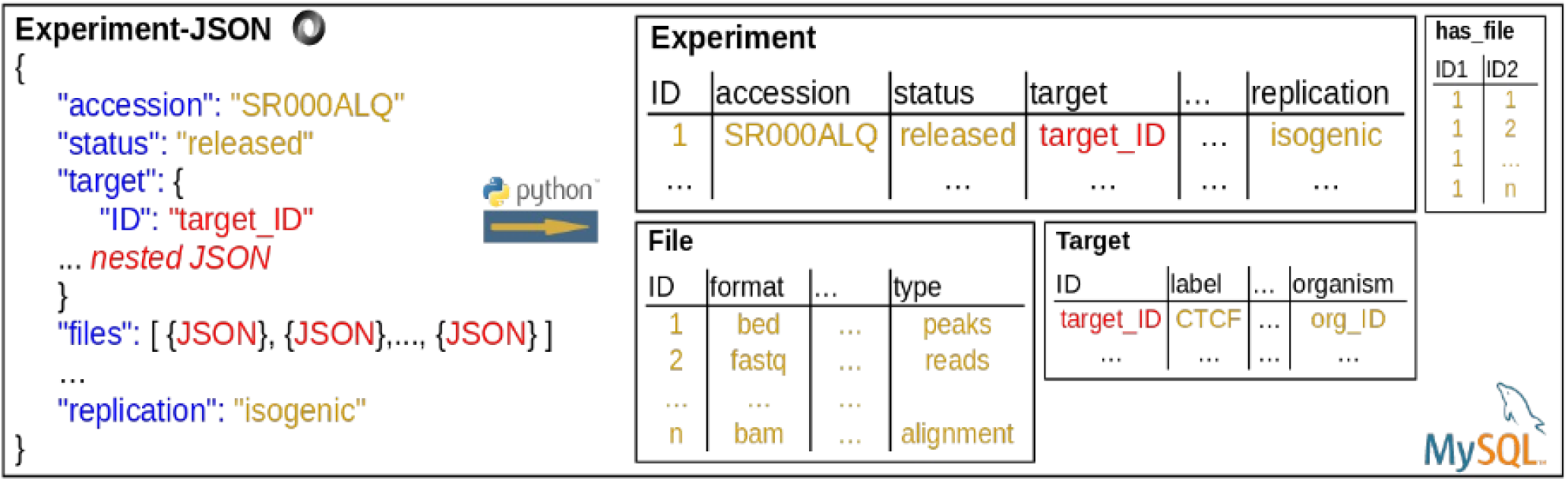
1) Same Cell Isogenic Line (K562) 2) Same condition/treatment 3) Same Laboratory (Snyder) 4) Same Bioinformatics ENCODE Pipeline 5) Same Statistical Test (IDR threshold)

**Figure S2.**
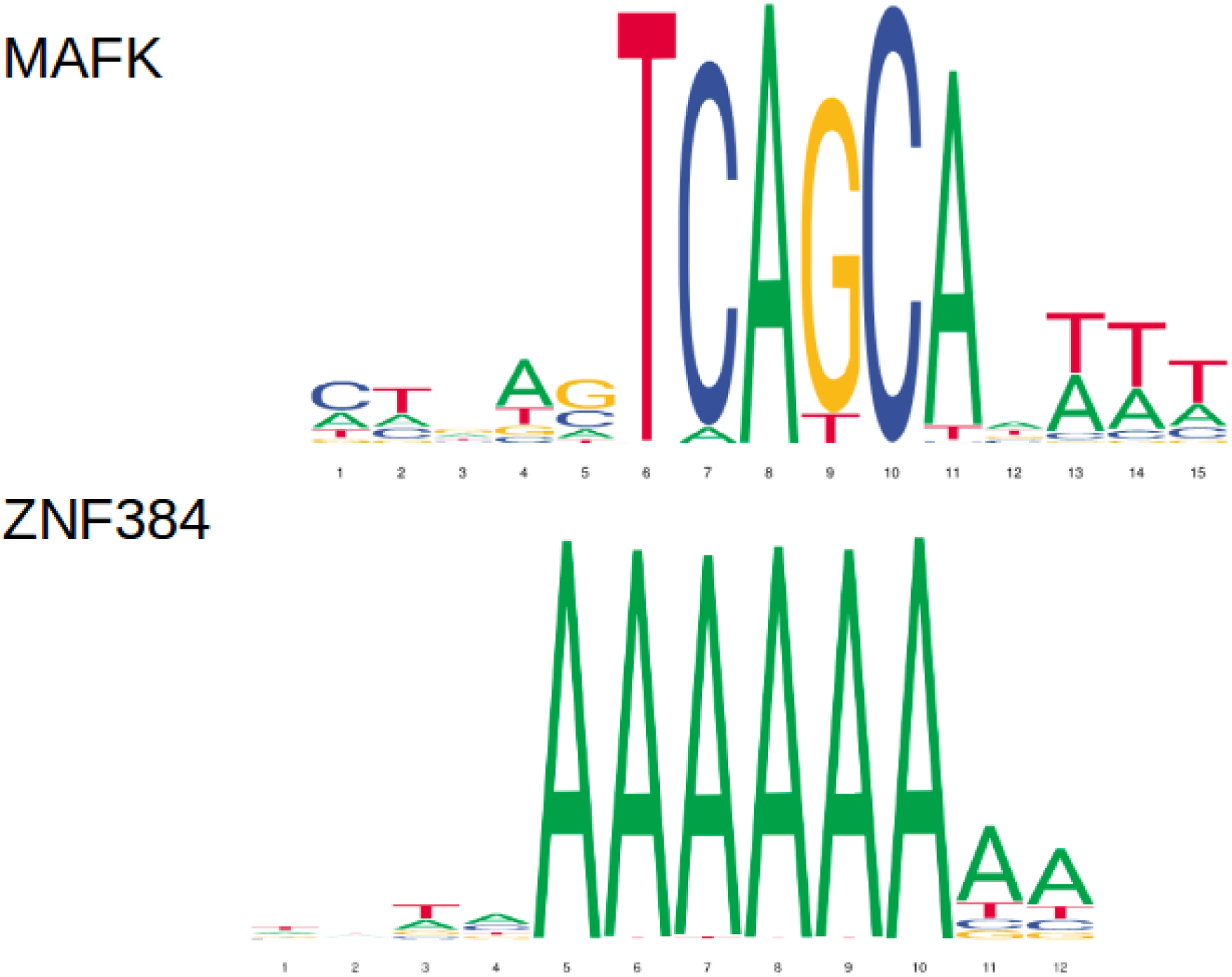
**(A)** DNA binding motifs of the protein targets of the ChIP-seq samples used to detect VOTs in mouse ESCs from the Jaspar database. No data for ZC3H11A and HCFC1.

**Figure S2.**
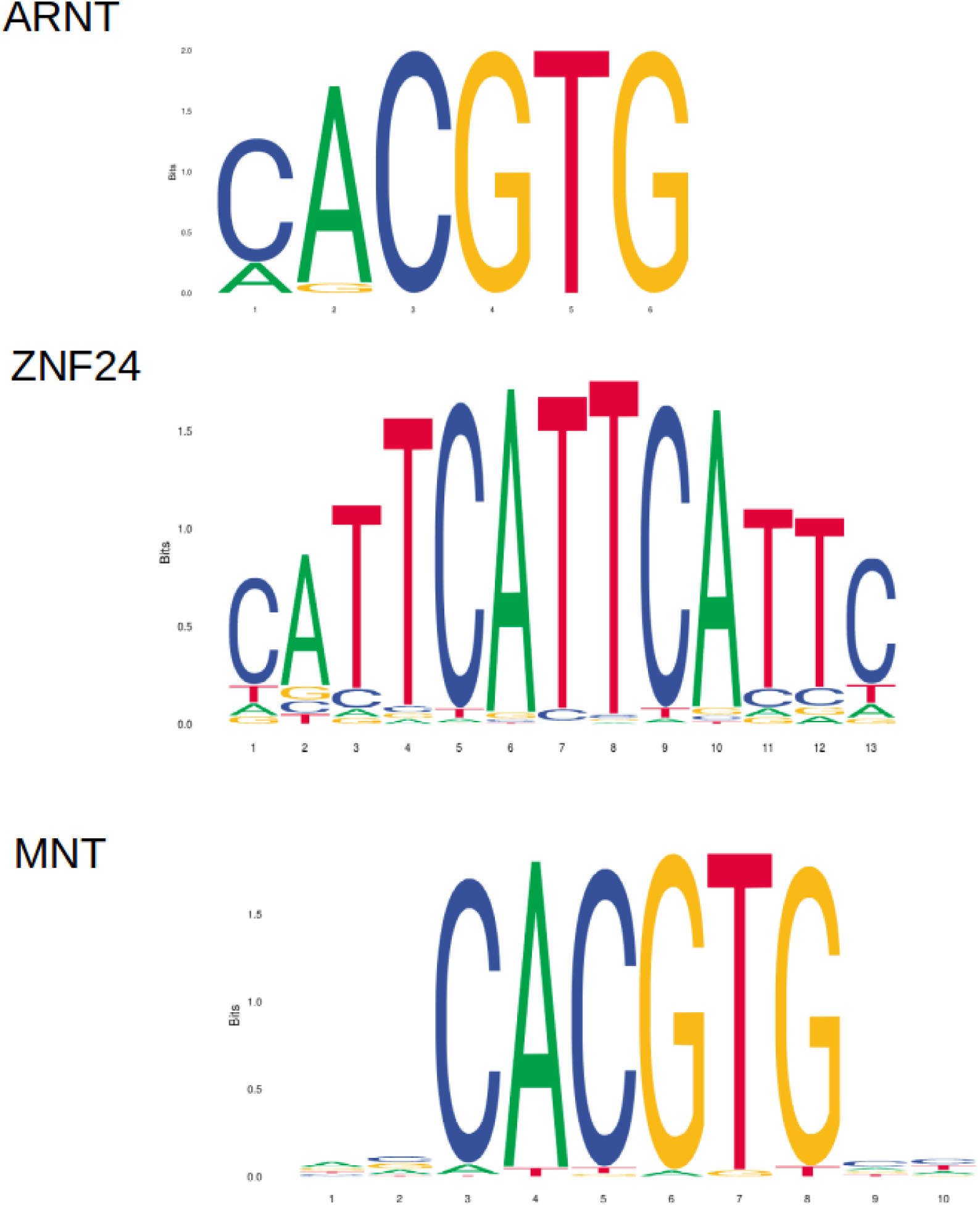
**(B)** DNA binding motifs of the protein targets of the ChIP-seq samples used to detect VOTs in K562 cell lines

**Figure S3.**
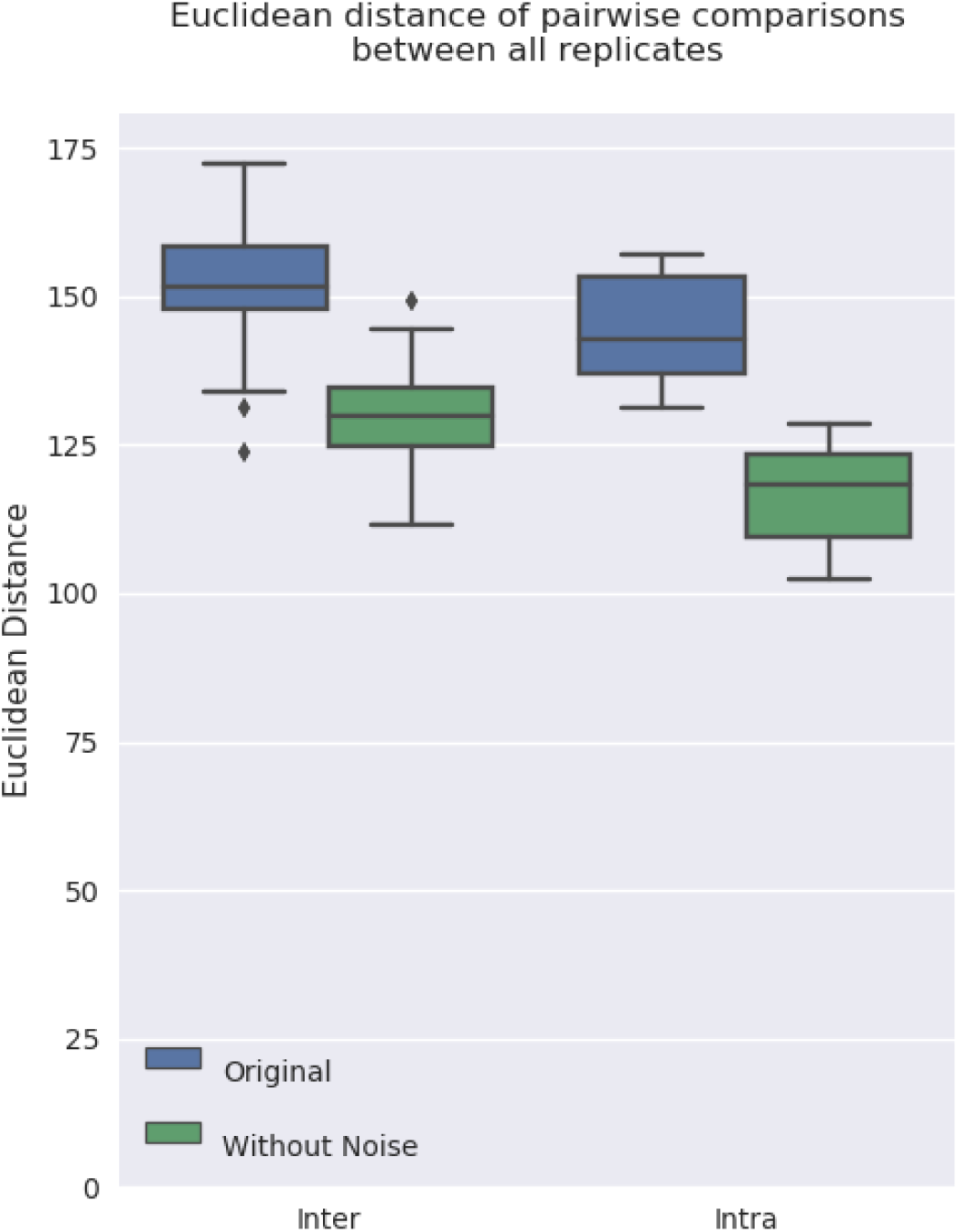
Pairwise comparisons of Euclidean distances between replicates of different proteins (inter) and within replicates of the same proteins (intra) in the original dataset and after removing the variable regions for the K562 ChIP-seq dataset (four proteins with three replicates).

